# Molecular Subtyping reveals Immune Alterations associated with Progression of Bronchial Premalignant Lesions

**DOI:** 10.1101/413898

**Authors:** Jennifer Beane, Sarah A. Mazzilli, Joshua D. Campbell, Grant Duclos, Kostyantyn Krysan, Christopher Moy, Catalina Perdomo, Michael Schaffer, Gang Liu, Sherry Zhang, Hangqio Liu, Jessica Vick, Samjot S. Dhillon, Suso J. Platero, Steven M. Dubinett, Christopher Stevenson, Mary E. Reid, Marc E. Lenburg, Avrum E. Spira

## Abstract

Bronchial premalignant lesions (PMLs) are precursors of lung squamous cell carcinoma, but have variable outcome, and we lack tools to identify and treat PMLs at highest risk for progression to invasive cancer. Profiling endobronchial biopsies of PMLs obtained from high-risk smokers by RNA-Seq identified four PML subtypes with differences in epithelial and immune processes. One molecular subtype (*Proliferative*) is enriched with dysplastic lesions and exhibits up-regulation of metabolic and cell cycle pathways and down-regulation of ciliary processes. RNA-Seq profiles from normal-appearing uninvolved large airway brushings could identify subjects with *Proliferative* lesions with high specificity. Expression of interferon signaling and antigen processing/presentation pathways are decreased in progressive/persistent *Proliferative* lesions and immunofluorescence indicates a depletion of innate and adaptive immune cells in these lesions. Molecular biomarkers measured in PMLs or the uninvolved airway can enhance histopathological grading and suggests that immunoprevention strategies may be effective in intercepting the progression of PMLs to lung cancer.

## Introduction

Lung cancer (LC) is the leading cause of cancer death taking about 160,000 U.S. lives each year, more than colorectal, pancreatic, breast, and prostate cancers combined. In order to decrease mortality, we need innovative strategies to intercept cancer development by diagnosing the disease at its earliest and potentially most curable stage. Recent advances based on results from the National Lung Screening Trial(1) are dramatically altering the landscape of early LC detection as computed tomography (CT) screening of high-risk individuals significantly reduces mortality. Despite this progress, biomarkers are needed to select individuals for LC screening as eligibility criteria account for less than 27% of individuals diagnosed with LC in the US(2) and to distinguish between benign or cancerous indeterminate pulmonary nodules as screening has very high false positive rate (>90%). There is also urgent and unmet need to develop personalized therapies earlier in the disease process to “intercept” LC prior to its development in this high-risk population.

Development LC risk biomarkers and LC interception strategies requires a detailed understanding of the earliest molecular alterations involved in lung carcinogenesis that occur in the respiratory epithelium*(3, 4)*. Exposure to cigarette smoke creates a field of injury throughout the entire respiratory tract by inducing a variety of genomic alterations that can lead to an “at-risk” airway where premalignant lesions (PMLs) and LCs develop. Lung squamous cell carcinoma (LUSC) arises in the epithelial layer of the bronchial airways and is often preceded by the development of a stepwise histological progression from normal epithelium to hyperplasia, squamous metaplasia, dysplasia (mild, moderate and severe), *carcinoma in situ* (*CIS*), and finally to invasive and then metastatic LUSC(5). In fact, the presence of high-grade persistent or progressive dysplasia (moderate or severe) is a marker of increased LC risk both at the lesion site (where they are the presumed precursors of squamous cell lung cancer) and elsewhere in the lung, although many dysplastic lesions do have varied outcomes*(6)*. Currently, however, we lack effective tools to identify PMLs at highest risk of progression to invasive carcinoma*(7)*. The development of markers of disease progression would identify patients at high-risk, suggest novel lung cancer chemoprevention agents, and provide molecular biomarkers for monitoring outcome in lung cancer prevention trials.

We hypothesize that molecular characterization of bronchial biopsies containing a mixture of epithelial and immune cells would allow us to identify transcriptomic alterations associated with high-grade histology and premalignant lesion progression. In this study, we used mRNA sequencing to profile endobronchial biopsies and brushings obtained through serial bronchoscopies from high-risk smokers undergoing lung cancer screening by auto-fluorescence bronchoscopy and chest CT. Using the bronchial biopsies, we identified four molecular subtypes associated with clinical phenotypes and biological processes. One subtype (*Proliferative* subtype) is enriched with biopsies having dysplastic histology, high basal cell and low ciliated cell signals, and expression of proliferation-associated pathways. Genes involved in interferon signaling and T cell mediated immunity were down-regulated among progressive/persistent lesions within the *Proliferative* subtype and these pathways correlated with decreases in both innate and adaptive immune cell types. Molecular classification of biopsies into a high-grade/progressive disease group may be used to stratify patients into prevention trials and to monitor efficacy of the treatment. The results also suggest that personalized lung cancer chemoprevention targeting specific cancer-related pathways or the immune system may have potential therapeutic benefits.

## Results

### Subject population

In this study, we used mRNA sequencing to profile endobronchial biopsies and brushings obtained through serial bronchoscopy of high-risk smokers undergoing lung cancer screening by auto-fluorescence bronchoscopy and CT at the Roswell Park Comprehensive Cancer Center (Roswell) in Buffalo, NY. The Discovery Cohort samples were obtained from the Roswell subjects between 2010 and 2012 (DC; n=29 patients, n=191 biopsies, n=91 brushes), and the Validation Cohort samples were obtained between 2012 and 2015 (VC; n=20 patients, n=111 biopsies, and 49 brushes). The subjects are predominantly older smokers, many of which have a history of lung cancer, chronic obstructive pulmonary disease (COPD), and occupational exposures that confer a high-risk of developing lung cancer. Clinical characteristics such as sex, age, smoking status (ever or never) reported at baseline visit, prior history of lung cancer, COPD status, and occupational exposures were not significantly different between the two cohorts (**Table 1**). After sample filtering based on several quality metrics, the DC had 190 biopsies and 89 brushes while the VC had 105 biopsies and 48 brushes. Ninety-four percent of subjects had at least one lung anatomic location sampled 2 or more times via endobronchial biopsy. The DC and VC contained 37.9% and 35.2% biopsies with a histological grade of dysplasia or higher and 23.1% and 19.0% had progressive/persistent dysplasia, respectively (**Table 2**). We used a previously described smoking-associated signature*(8)* to predict the smoking status of each sample, as smoking status was only available at baseline, and found that the DC had a higher percentage of biopsies predicted to be current smokers (62.6%) compared with the VC (36.2%). There is no significant difference in smoking status among the bronchial brushings between the two cohorts since only 1 brush is collected per time point. In terms of RNA sequencing quality, the DC had significantly greater total reads, percent uniquely mapping reads, and median transcript integrity number scores among the biopsies than the VC, but these differences between cohorts were not reflected in the brushes (**Table S1**).

**Table 1.** Demographic and Clinical Annotation on Subjects in both the Discovery and Validation cohorts. Statistical tests between the Discovery and Validation cohorts were performed using Fisher’s Exact Test for categorical variables and Student’s T-Test for continuous variable. Percentages are reported for categorical variables and mean and standard deviations are reported for continuous variables.

| Variable | Discovery Cohort<br>(n=30 Subjects) | Validation Cohort<br>(n=20 Subjects) | p-value |
| --- | --- | --- | --- |
| Average # Biopsies/Subject | 6.6 (5.7) | 5.25 (2.9) | 0.3 |
| Average # Bronchoscopies/Subject | 2.8 (1.5) | 2.4 (0.8) | 0.27 |
| Average Time Between Bronchoscopies (Days) | 368.2 (201.4) | 360.1 (212.5) | 0.87 |
| Male | 15/30 (50) | 12/20 (60) | 0.57 |
| White | 27/30 (90) | 17/20 (85) | 0.67 |
| Age (at Baseline Clinical Visit) | 58.8 (7.6) | 58.7 (8.3) | 0.97 |
| Ever smoker (at Baseline Clinical Visit) | 29/30 (96.7) | 19/20 (95) | 1 |
| Prior History of Lung Cancer | 21/30 (70) | 12/20 (60) | 0.55 |
| COPD (FEV1/FVC <= 0.7, at Baseline Clinical Visit) | 17/27 (63.0) | 8/18 (44.4) | 0.24 |
| GOLD 1 (FEV1% > 80) | 2/27 (7.4) | 2/18 (11.1) | 1 |
| GOLD 2 (FEV1% <80 and > 50) | 12/27 (44.4) | 5/18 (27.8) | 0.35 |
| GOLD 3 (FEV1% < 50 and > 30) | 3/27 (11.1) | 1/18 (5.6) | 0.64 |
| Occupational Asbestos | 13/30 (43.3) | 9/20 (45) | 1 |
| Occupational High-Risk Job | 14/30 (46.7) | 12/20 (60) | 0.4 |

**Table 2.** Clinical Annotation on Samples in both the Discovery and Validation cohorts. Statistical tests between the Discovery and Validation cohorts within either the biopsies or brushes were performed using Fisher’s Exact Test and percentages are reported.

| Variable | Discovery Cohort |  | Validation Cohort |  | P-value |  |
| --- | --- | --- | --- | --- | --- | --- |
| Sample Type | Biopsies | Brushes | Biopsies | Brushes | Biopsies | Brushes |
| Histology |  |  |  |  | 0.05 | 0.42 |
| Normal | 38/190 (20) | 6/89 (6.7) | 23/105 (21.9) | 0/48 (0) |  |  |
| Hyperplasia | 30/190 (15.8) | 11/89 (12.4) | 31/105 (29.5) | 9/48 (18.8) |  |  |
| Metaplasia | 46/190 (24.2) | 15/89 (16.9) | 14/105 (13.3) | 9/48 (18.8) |  |  |
| Mild Dysplasia | 21/190 (11.1) | 9/89 (10.1) | 13/105 (12.4) | 6/48 (12.5) |  |  |
| Moderate Dysplasia | 38/190 (20) | 30/89 (33.7) | 20/105 (19.0) | 18/48 (37.5) |  |  |
| Severe Dysplasia | 12/190(6.3) | 17/89 (19.1) | 4/105 (3.8) | 6/48 (12.5) |  |  |
| CIS | 1/190 (0.5) | 0/89 (0) | 0/105 (0) | 0/48 (0) |  |  |
| Tumor | 0/190 (0) | 1/89 (1.1) | 0/105 (0) | 0/48 (0) |  |  |
| Unknown Histology | 4/190 (2.1) | 0/89 (0) | 0/105 (0) | 0/48 (0) |  |  |
| Current smoker (Genomic prediction) | 119/190 (62.6) | 44/89 (49.4) | 38/105 (36.2) | 20/48 (41.7) | 1.80E-05 | 0.47 |
| Progression Status |  |  |  |  | 0.39 |  |
| Normal/Stable | 47/190 (24.7) |  | 35/105 (33.3) |  |  |  |
| Progressive/Persistent | 44/190(23.2) |  | 20/105 (19.0) |  |  |  |
| Regressive | 30/190 (15.8) |  | 18/105 (17.1) |  |  |  |
| Unknown | 69/190 (36.3) |  | 32/105 (30.5) |  |  |  |

### LUSC PMLs within the discovery cohort divide into distinct molecular subtypes

In order to identify gene expression differences associated with LUSC PML histological severity using the endobronchial biopsies, we used a discovery-based approach to identify *de novo* molecular subtypes based on distinct patterns of gene co-expression (gene modules). The approach was chosen given that there is histological heterogeneity within biopsies and that pathological analyses were conducted using biopsies adjacent to biopsies profiled via mRNA-Seq. First, we sought to select a set of gene modules that are present across different LUSC datasets. Using weighted gene co-expression network analysis(9) (WGCNA), gene modules were derived in the DC biopsies (n=190 samples, n=16653 genes, n=15 gene modules), the DC brushes (n=89 samples, n=16058 genes, n=47 gene modules), TCGA squamous cell carcinoma (LUSC) tumors(10) (n=471 samples, n=17887 genes, n=55 gene modules), and tracheobronchial samples from mice treated with n-nitrosotris-(2-choroethyl)urea (NTCU) (n=25 samples, n=14897 genes, n=40 gene modules). DC biopsy gene modules that were highly correlated (r>0.85) to at least one other non-DC biopsy module within each of the 4 datasets were selected. Genes in the selected modules were filtered by requiring that each gene was also present in at least one of the correlated non-DC biopsy modules, resulting in a set of 9 gene modules that consisted of 3,936 genes in total (**Table S2)**. These gene modules identified 4 molecular subtypes within the DC biopsies via consensus clustering: *Proliferative* (dark blue, n=52 samples, 27.4%), *Inflammatory* (dark green, n=37 samples, 19.5%), *Secretory* (light blue, n=61 samples, 32.1%), and *Normal* (light green, n=40 samples, 21.1%) (**Fig. 1A, Table 3**).

**Table 3.**
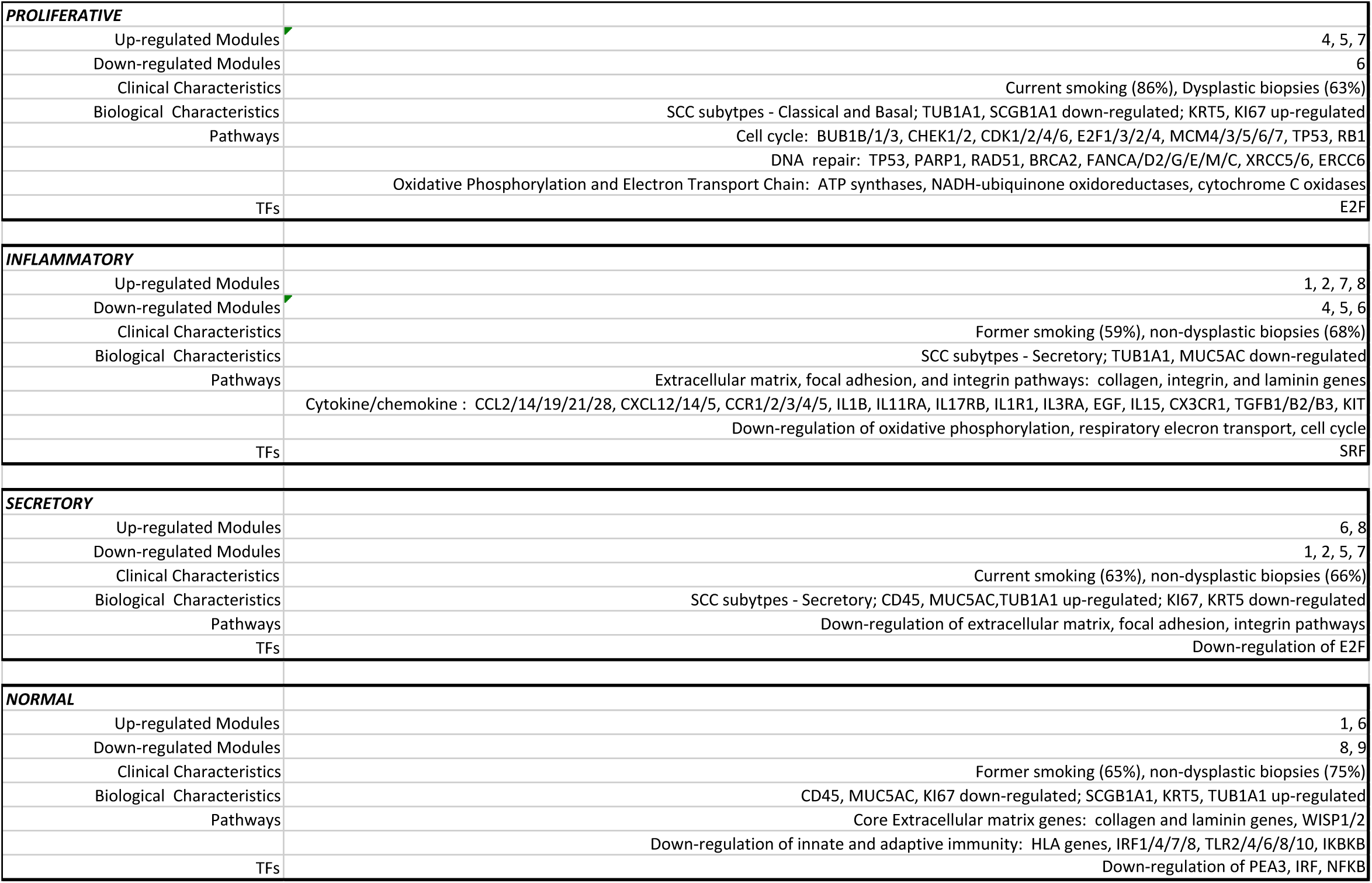
Summary of Molecular Subtype Characteristics in the Discovery Cohort. For each molecular subtype, significant associations are reported with each of the 9 gene modules, clinical characteristics, canonical cell type epithelial and white blood cell gene markers, pathways, and transcription factors.

**Figure 1.**
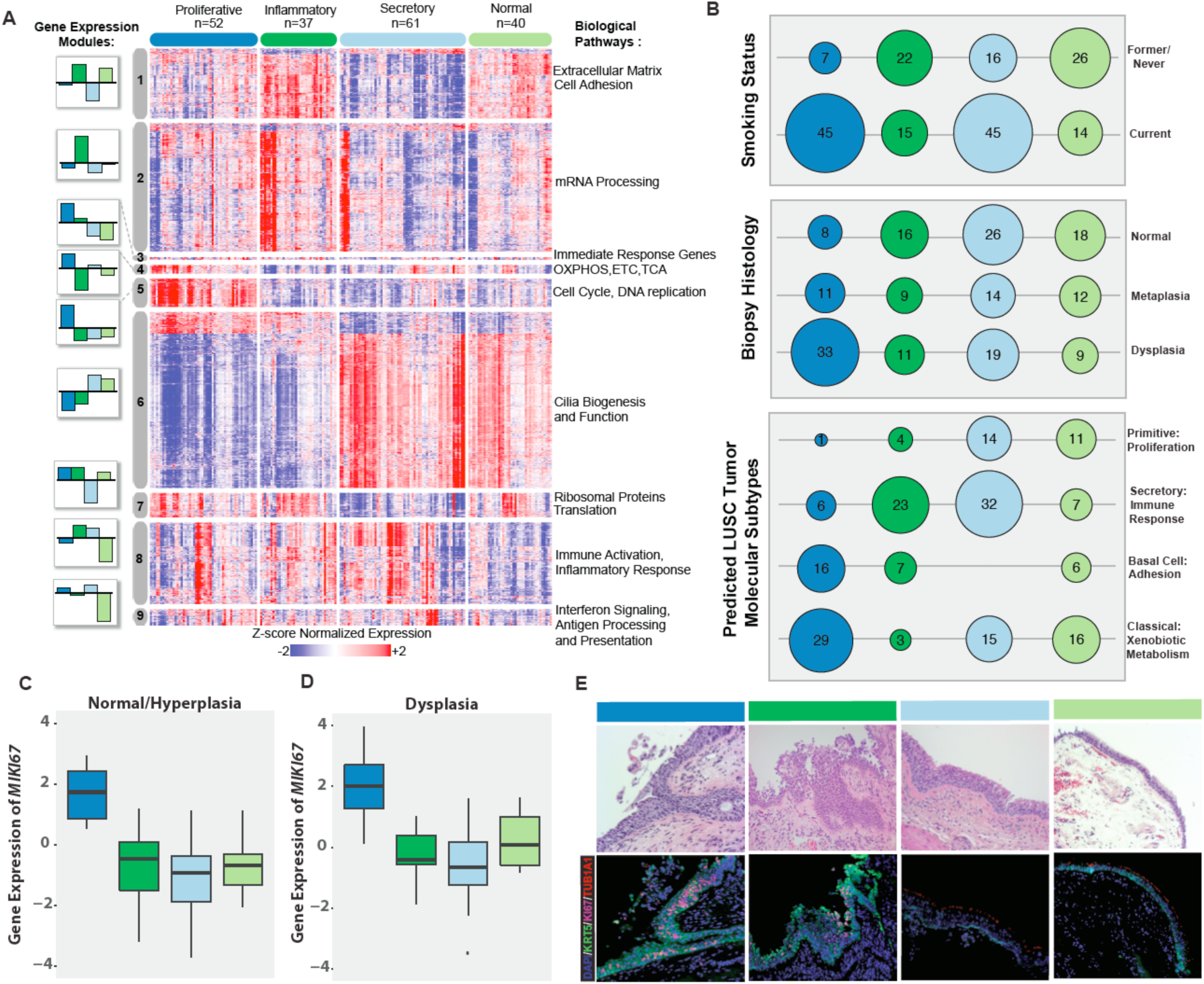
Endobronchial biopsies divide into four distinct molecular subtypes that correlate with clinical and molecular phenotypes. (**A**) Genes (n=3,936) organized into 9 gene co-expression modules were used to discover four molecular subtypes (*Proliferative, Inflammatory, Secretory*, and *Normal*) across the 190 DC biopsies using consensus clustering. The heatmap shows semi-supervised hierarchal clustering of z-score normalized gene expression across the 3,936 genes and 190 DC biopsies. The top color bar represents the four molecular subtypes: *Proliferative* (n=52 samples), *Inflammatory* (n=37 samples), *Secretory* (n=61 samples), and *Normal* (n=40 samples). On the left side of the heatmap, the mean of the first principal component calculated across module genes is plotted for each subtype. On the right side of the heatmap, a summary of enriched biological pathways is listed for each module. (**B**) Bubbleplots showing significant associations (p < 0.01 by Fisher’s Exact Test) between the molecular subtypes and smoking status, biopsy histological grade, and the predicted LUSC tumor molecular subtypes. The columns represent the 4 molecular subtypes (*Proliferative, Inflammatory, Secretory*, and *Normal*) and the diameter of the circle is proportional to the number of samples within each subtype that have the row phenotype. (**C**) Boxplot of expression values of *MKI67* in biopsies with normal or hyperplasia histology (n=8, 16, 26, 18 in *Proliferative, Inflammatory, Secretory*, and *Normal* subtypes, respectively). The *MKI67* expression levels of the *Proliferative* subtype are significantly greater than non-*Proliferative* subtype samples (FDR=3.4e-10) (**D**) Boxplot of expression values of *MKI67* in biopsies with dysplastic histology (n=33, 11, 19, 9 in *Proliferative, Inflammatory, Secretory*, and *Normal* subtypes, respectively). The *MKI67* expression levels of the *Proliferative* subtype are significantly greater than non-*Proliferative* subtype samples (FDR=3.1e-8). (**E**) Immunofluorescent staining demonstrating the increased *MKI67* and *KRT5* staining and reduced *TUB1A1* staining in the Proliferative subtype in concordance with the expression of the corresponding marker genes. The representative samples shown for the Proliferative and Inflammatory subtypes have dysplasia histology while the samples shown for the Secretory and Normal subtypes have normal histology (Magnification 200X).

In order to characterize each molecular subtype, we first focused on identifying biological pathways over-represented in the genes comprising each gene module, as the pattern of gene module expression defines each PML subtype. Each gene module was found to be associated with distinct epithelial and immune biological processes (**Fig. 1A and Tables S2 and S3**). The *Proliferative* subtype is specifically characterized by increased expression of genes involved in energy metabolism and cell cycle pathways (Modules 4 and 5). The *Secretory* and *Normal* subtypes both have increased expression of genes in cilium-associated pathways (Module 6), however, the *Normal* subtype specifically has decreased expression of genes involved in inflammation, regulation of lymphocytes and leukocytes, and antigen processing and presentation pathways (Modules 8 and 9). The *Secretory* subtype exhibits decreased expression of genes involved in protein translation (Module 7), while RNA processing genes (Module 2) are expressed more highly in the *Inflammatory* subtype.

We further characterized our molecular subtypes by their associations with clinical phenotypes and established LUSC tumor molecular subtypes*(11, 12)*. Sample smoking status, the subject from whom the sample was derived, and sample histology demonstrated significant associations with subtype (p<0.01, **Fig. 1B, Table S4, Fig. S1**). Our *Proliferative* and *Secretory* subtypes are enriched for current smokers and this association drives the subject enrichment as 79% of subjects maintain their smoking status throughout the study. Additionally, the *Proliferative* subtype is enriched for biopsies with dysplasia histology (**Fig. 1B**). The *Proliferative* subtype has high expression of genes involved in cell cycle processes including the proliferation marker *MIKI67,* which is significantly up-regulated among samples in this subtype compared with samples in other subtypes (FDR=1.0e-30, based on differential expression analysis between samples in the *Proliferative* versus the non-*Proliferative* subtypes across all genes). The gene remained significantly up-regulated in the *Proliferative* subtype within samples with normal/hyperplasia histology (FDR=3.4e-10) and samples with dysplasia histology (FDR=3.1e-8), and these observations are supported by an increase in protein expression in representative samples (p=0.02) (**Fig. 1C-E and Fig. S2**). The *Proliferative* subtype samples also had high concordance with the LUSC-Classical subtype (**Fig. 1B**). In the TCGA LUSC tumors, the LUSC-Classical subtype was associated with alterations and overexpression of *KEAP1* and *NFE2L2* as well as amplification of 3q26 with overexpression of *SOX2, TP63* and *PIK3CA*(11). Similarly, our *Proliferative* PMLs have increased expression of *KEAP1, NFE2L2, TP63*, and *PIK3CA* (FDR=1.4e-6, 4.5e-12, 1.4e-9, and 0.03, respectively) (**Fig. S3A**). Furthermore, the LUSC-Classical subtype was found to be associated with increased expression of genes involved in energy metabolism, and our *Proliferative* subtype is in part defined by high expression of Module 4, which is enriched for genes associated with oxidative phosphorylation and the electron transport chain. In contrast, the *Inflammatory* and *Secretory* PML subtypes demonstrate enrichment for the LUSC-Secretory subtype. The LUSC-Secretory subtype was associated with processes related to the immune response, and the *Inflammatory* and *Secretory* PMLs have the highest expression of Module 8 that is enriched for genes in these same pathways.

Finally, we wanted to examine the extent to which our PML molecular subtypes were driven by differences in epithelial and immune cell type composition by assessing expression of a number of canonical cell type markers. The *Inflammatory* and *Secretory* subtypes have higher levels of expression of the white blood cell marker *PTPRC (CD45)* consistent with enrichment of the LUSC-Secretory subtype (**Fig. S3B,** FDR=0.12 and 0.01, respectively). Consistent with the behavior and pathways enriched in Module 6, the ciliated cell marker *TUB1A1* expression is decreased in the *Inflammatory* and *Proliferative* subtypes (FDR=1.1e-4 and 3.5e-19, respectively), and this is also shown by a decrease in acetylated a-tubulin staining in representative histological samples (**Fig. 1E, Fig. S2**). The *Proliferative* subtype has the highest expression (FDR=2.4e-15) of basal cell marker *(KRT5*) indicating enrichment of lesions with high-grade histology that tightly correlates with protein expression in representative histology samples (p=0.01) (**Fig. 1E, Fig. S2, Fig. S3B, Table S5**). Additionally, gene expression of *MUC5AC*, a marker of goblet epithelial cells, is increased in subtypes enriched for current smokers (*Proliferative* and *Secretory*) but is the most significantly increased in the *Secretory* subtype (FDR=3.4e-5). In contrast, gene expression of *SCGB1A1,* a marker of club cells, is the lowest in the *Proliferative* subtype (FDR=6.1e-5). The expression levels of these marker genes agree with cell type deconvolution methods to examine epithelial and immune cell content (**Fig. S3C-D**). The summation of these characterizations highlights epithelial and immune cell associated pathways that are modulated by smoking and PML histology and identifies the *Proliferative* subtype as a subset of high-grade PMLs that express proliferative and cell cycle-related pathways.

### Phenotypic associations with the molecular subtypes are confirmed in the Validation Cohort

Next, we wanted to determine if the heterogeneity captured in the DC biopsy-derived molecular subtypes was reproducible in the VC. We developed a 22-gene nearest centroid molecular subtype predictor by selecting genes representative of each of the 9 gene modules. The predictor has 84.7% accuracy across DC biopsies (training set, **Fig. 2A** and **Fig. S4)** with the following misclassification rates per subtype 5/52 (9.6%) in *Proliferative*, 7/37 (18.9%) in *Inflammatory*, 9/61 (14.8%) in *Secretory*, and 8/40 (20%) in *Normal*. The 22-gene classifier was used to predict the molecular subtype of the 105 VC biopsies (**Fig. 2B**). The VC subtype predictions were evaluated by examining the concordance of metagene scores for each of the 9 modules (using the full set of genes for each module) between the predicted VC subtypes compared with the DC subtypes. The average behavior of Principal Component 1 (PC1) across the subtypes was highly similar (**Fig. S5**) with few exceptions (namely, Module 3 that had the fewest genes). Additionally, we compared the VC subtype predictions from the 22-gene classifier to subtypes derived in the VC biopsies using the same methodology used to derive the DC subtypes and found significant concordance (p=1.0e-7, with the *Proliferative* subtype having the greatest concordance between predictions, **Fig. S4).**

**Figure 2.**
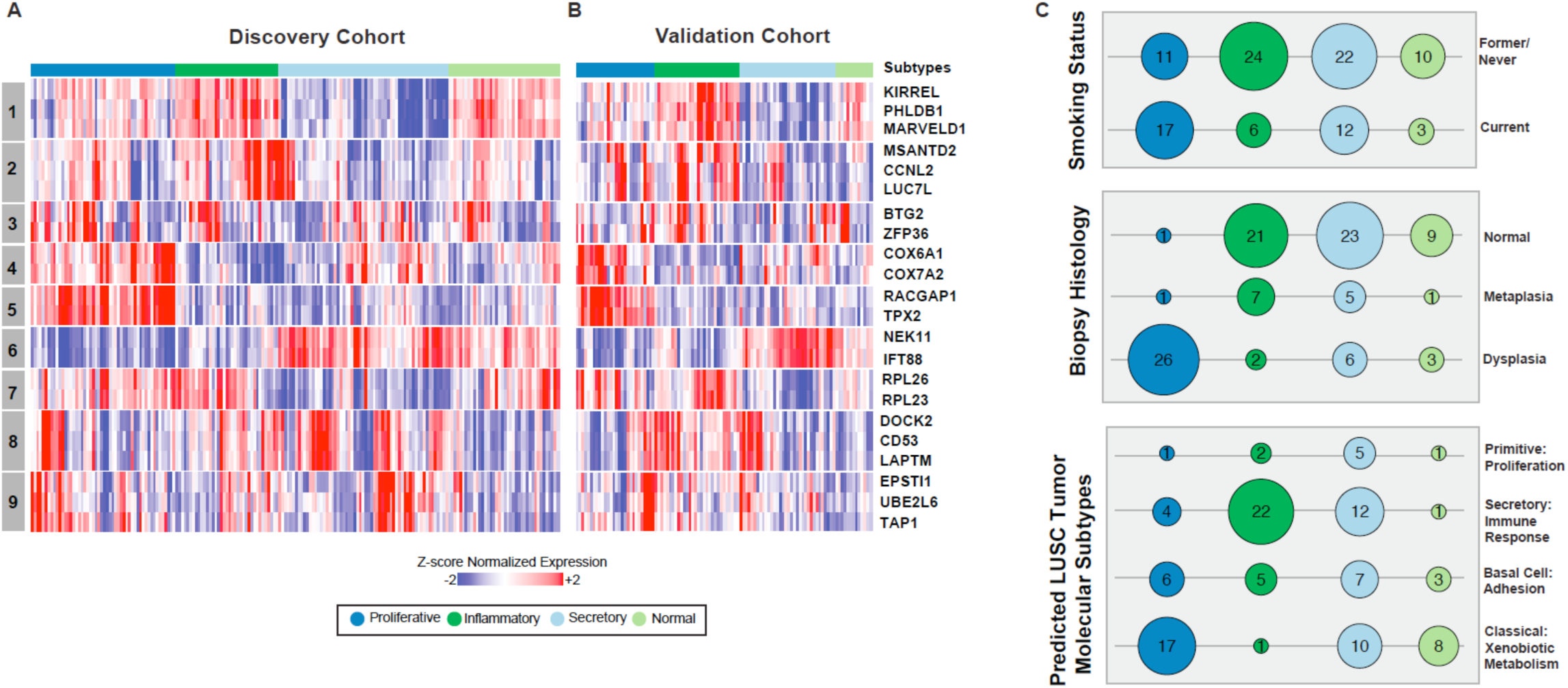
Phenotypic associations with the molecular subtypes are confirmed in an independent sample set. (**A**) The 190 DC biopsies and the 3,936 genes were used to build a 22-gene nearest centroid molecular subtype classifier. Semi-supervised hierarchal clustering of z-score normalized gene expression across the 22 classifier genes and 190 DC biopsies training samples. (**B**) The 22-gene nearest centroid molecular subtype classifier was used to predict the molecular subtypes of the 105 VC biopsies. Semi-supervised hierarchal clustering of z-score normalized gene expression across 22 genes and 105 VC is plotted. The rows of the heatmap give the gene name and module membership, and the column color bar shows molecular subtype membership. (**C**) Bubbleplots showing significant associations (p<0.01 by Fisher’s Exact Test) between the VC molecular subtypes and smoking status, biopsy histological grade, and the predicted LUSC tumor molecular subtypes. The columns represent the 4 molecular subtypes (*Proliferative, Inflammatory, Secretory*, and *Normal*) and the radius of the circle is proportional to the number of samples within each subtype that have the row phenotype.

The statistical associations between the VC subtypes (via the 22-gene classifier) and clinical and molecular phenotypes across the VC biopsies are analogous to those observed across the DC biopsies (**Fig. 2C, Table S4, Fig. S1 and S3**). Briefly, the *Proliferative* subtype is enriched for current smokers, biopsies with dysplasia histology, and the LUSC-Classical tumor subtype (**Fig. 2C, Table S4**). Epithelial and white blood cell marker gene expression across the VC biopsies reveals higher levels of the white blood cell marker *PTPRC* (*CD45* expression) in the *Inflammatory* subtype (FDR=0.002) consistent with enrichment of the LUSC-Secretory subtype (**Fig. S3F)**. The *Inflammatory* and *Proliferative* subtypes have reduced ciliated cell marker expression (*FOXJ1*) consistent with Module 6 (*FOXJ1* FDR=0.0005 and FDR=2.62e-6 and Module 6 FDR=5.73e-6 and FDR=4.34e-10, respectively). The *Proliferative* subtype has the highest expression of basal cell marker *KRT5* (FDR=1.67e-7), proliferation marker *MKI67* (FDR=3.03e-10), and cell cycle associated Module 5 (FDR=1.23e-18) indicating enrichment of lesions expressing characteristics associated with high-grade histology. Gene expression of *SCGB1A1,* a marker of club cells, is the lowest in the *Proliferative* subtype (FDR=1.8e-4). Gene expression of *MUC5AC*, a marker of goblet epithelial cells, was increased in current smokers and most significantly in the *Secretory* subtype in the DC biopsies; however, in the VC biopsies this trend is not preserved as current smokers are not enriched in the *Secretory* subtype. The expression levels of these marker genes agree with other deconvolution methods to examine epithelial and immune cell content (**Fig. S3E-H**).

### Normal appearing airway field brushes reflect biopsy molecular subtype

Previously, we have shown that bronchial brushes from normal appearing areas of the mainstem bronchus could predict the presence of PMLs(13); however, that study lacked biopsies and brushes from the same subjects. Above, in both the DC and the VC biopsies, the *Proliferative* subtype, represents a distinct subtype of PMLs enriched for dysplastic histology expressing metabolic and proliferative pathways. Biopsies classified as the *Proliferative* subtype may represent a group of PMLs that need close monitoring and intervention. As a result, we sought to explore whether or not we could predict the presence of *Proliferative* subtype biopsies using the brushes. The *Proliferative* subtype is defined by the behavior of Modules 4, 5, 6, and 7 (**Table 3**), and therefore, we used the subset of 8 genes (from the 22-gene predictor) that correspond to these Modules to predict the presence of the *Proliferative* subtype across the DC and VC biopsies and brushes. A prediction of the *Proliferative* subtype in a brush is specific (91% and 92% in the DC and VC biopsies, respectively), but not sensitive (39% and 32% DC and VC biopsies, respectively) at indicating the presence of at least one *Proliferative* PML detected at the same time point (**Fig. 3A**). In order to understand the classifier’s performance in predicting the *Proliferative* subtype in brushes, we examined Gene Set Variation Analysis (GSVA)(14) scores for Modules 4, 5, 6, and 7 that define the *Proliferative* subtype in the DC and VC brushes (**Fig. 3B**). In the DC and VC brushes, the GSVA scores were significantly different (FDR<0.05) in the *Proliferative* subtype versus all other samples only for Modules 5 and 6, and thus these likely contribute the most heavily to *Proliferative* subtype classification in the brushes. Module 5 contains genes associated with cell cycle and proliferation while Module 6 contains genes associated with cilium assembly and organization. Down-regulation of Modules 5 and 6 in the brushes specifically predicts the presence of a *Proliferative* subtype PML; however, the absence of these signals in the airway field of injury does not preclude the development of a *Proliferative* subtype PML.

**Figure 3.**
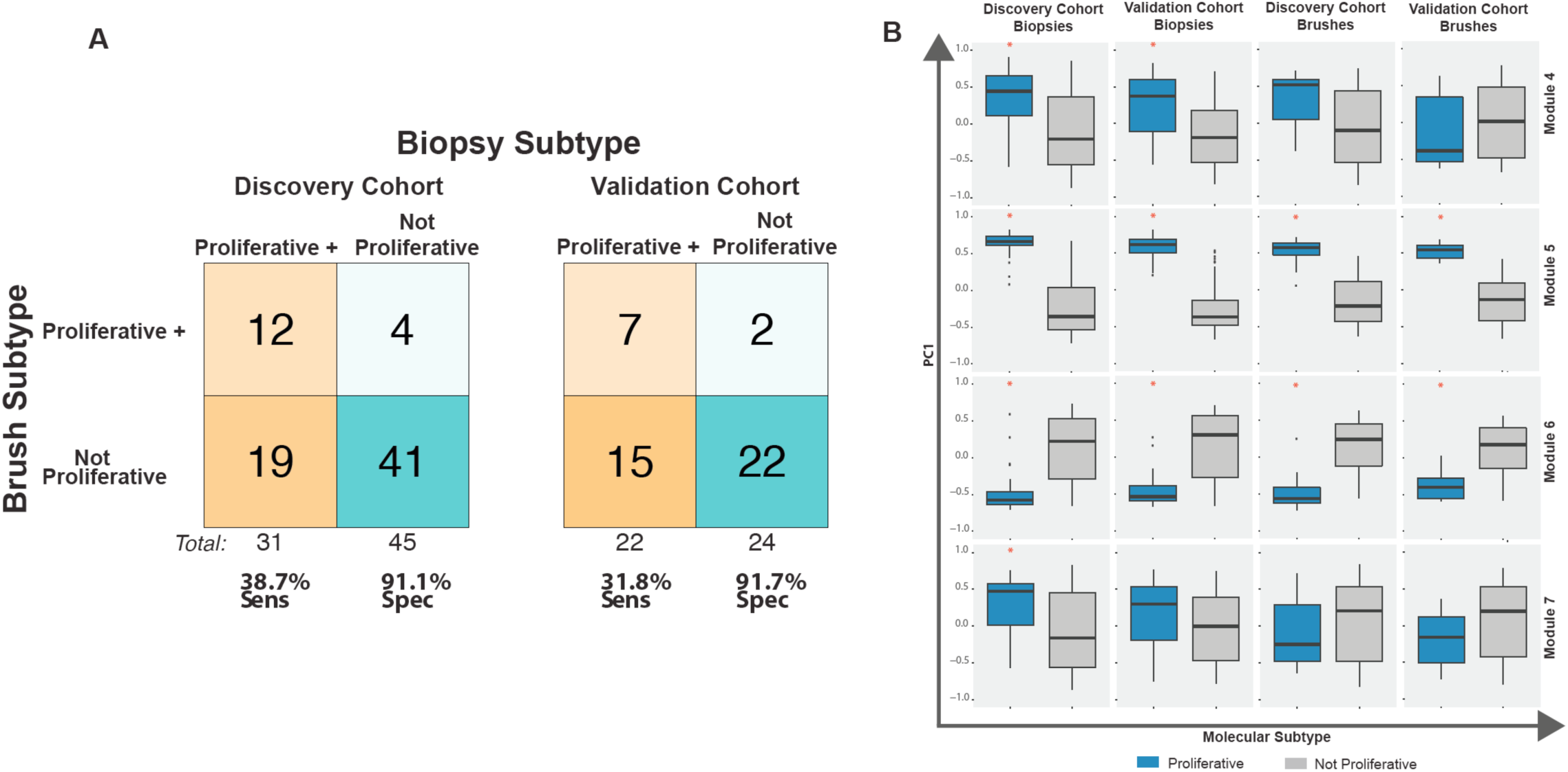
Performance of the molecular subtype classifier in the large airway brushes from normal appearing epithelium sampled at the same time as the endobronchial biopsies. (**A**) The DC (left) and VC (right) cohorts, showing the number of brushes (y-axis) predicted to be positive for the *Proliferative* subtype (orange) that have at least one biopsy (y-axis) with a classification of the *Proliferative* subtype at the time the brush was sampled. Brushes/biopsies negative for the *Proliferative* subtype are turquoise. (**B**) Boxplots of PC1 for Modules 4, 5, 6, and 7 (y-axis) across the four molecular subtypes for each cohort (x-axis). The red asterisk indicates significant differences between the *Proliferative* subtype versus all other samples (FDR<0.05).

### Immune-associated genes separate proliferative subtype progressive/persistent and regressive PMLs

Previous studies of bronchial PMLs suggest that high-grade lesions (which occur more frequently in current smokers) are more likely to progress to invasive carcinoma(6). Therefore, we sought to identify molecular alterations associated with subsequent PML progression/persistence (n=15) versus regression (n=15) among the *Proliferative* subtype DC biopsies, as these may be clinically relevant to identifying appropriate interception strategies. Using GSVA scores calculated across all the DC biopsies for each of the 9 modules, we calculated which scores were statistically different between progressive/persistent versus regressive disease in the samples belonging to the *Proliferative* subtype (**Table S6**). We found that the DC biopsy GSVA Module scores for Module 9 were significantly higher among regressive *Proliferative* PMLs (p=0.002, **Fig. 4A**) compared with progressive/persistent *Proliferative* PMLs. The association between low Module 9 score and progression/persistence is replicated in the VC biopsies (n=7 progressive/persistent and n=13 regressive biopsies; p=0.03, **Fig. 4B**). The ability of the Module 9 GSVA scores to discriminate between regressive versus progressing/persistent biopsies as measured by the area under the receiver operating characteristic (ROC) was 0.809 and 0.802 in the DC and VC biopsies, respectively.

**Figure 4.**
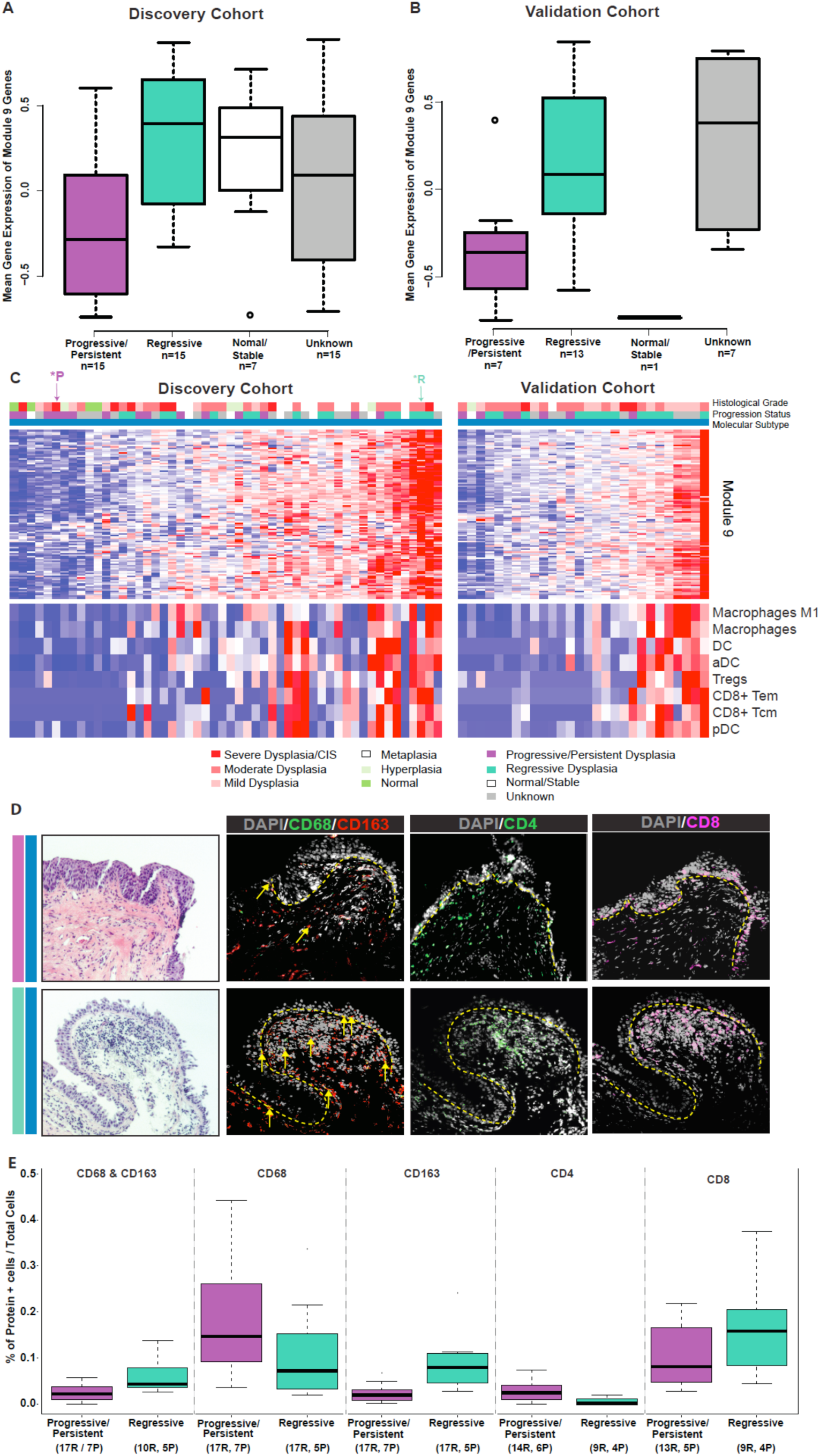
The module enriched for interferon signaling and antigen processing is associated with biopsy progression/persistence and a depletion of innate and adaptive immune cells in the Proliferative subtype. (**A**) Metagene expression of Module 9 genes among DC biopsies within the *Proliferative* subtype (p=0.002 between the progressive/persistent versus regressive biopsies). Biopsy progression/regression was defined for each biopsy based on the histology of the biopsy and the worst histology recorded for the same lung anatomic location in the future. Histology changes between normal, hyperplasia, and metaplasia were classified as “normal stable”, decreases in histological dysplasia grade or changes from dysplastic histology to normal/hyperplasia/metaplasia were classified as “regressive”, lack of future histological data was classified as “unknown”, and everything else was classified as “progressive/persistent.” (**B**) Metagene expression of Module 9 genes among VC biopsies within the *Proliferative* subtype (p=0.03 between the progressive/persistent versus regressive biopsies). (**C**) *Top:* Z-score normalized gene expression across the 112 genes in Module 9 and the DC biopsies (left) and the VC biopsies (right). Each heatmap is supervised according to the Module 9 GSVA scores. Top color bars indicate the histological grade of the biopsies and their progression status. *Bottom:* xCell results indicating the relative abundance of immune cell types across the DC biopsies (left) and the VC biopsies (right). Immune cell types displayed are significantly associated with lesion progressive/persistence (FDR<0.05 in both the DC and VC after adjusting for differences in epithelial cell content). (**D**) Representative histology where the dashed yellow line denoted the separate of epithelium and stromal compartment *Top panels*: A progressive severe dysplasia has reduced presence of immune cells demonstrated by the marked reduction in expression of M2 macrophages (CD68/163 staining, double positive cells indicated by the yellow arrows) and CD8 T cells. (sample corresponds to *\*P* in panel C.) *Bottom panels*: A regressive moderate dysplasia has increased presence of immune cells including M2 macrophages (CD68/163 staining double positive cells indicated by the yellow arrows) and CD8 T cells. (samples correspond to *\*R* in panel C.) (**E**) Boxplots of the percentages of CD68 and CD163, CD68, CD163, CD4, and CD8 positively stained cells between progressive/persistent and regressive biopsies (p<0.001 for all comparisons). The x-axis labels indicate the number of regions (R) enumerated across (P) subjects for each stain and outcome group depicted in the boxplot. Biopsies were included in the analysis if their clinical outcome was concordant with the Module 9 score.

The genes in Module 9 include a number of genes that encode for proteins involved in interferon signaling as well as antigen processing and presentation (*SP100, CIITA, CXCL10, SOCS1, GBP1, GBP4, B2M, TAP1, TAPBP, TRIM 14, TRIM21, TRIM22, STAT1, PML, OAS2, OAS3, MX1, ADAR, ISG15, IFI35, IFIT3, IFI27, PSMB8, PSMB9, BST2, IRF1, IRF9, CD74, PSME1, PSME2, HLA-DQA1/DPA1/ DPB1/DRA/ DQB2/DRB1/ DQB1/DMA/DMB/DOA, HLA-A/B/C/E/F*) and include the inhibitory receptor *LAG3*. As a result, we wanted to evaluate whether or not the presence or absence of innate or adaptive immune cells were associated with Module 9 expression within the Proliferative subtype. In an effort to deconvolute the potential presence of immune cell types, we generated GSVA scores using previously described immune cell signatures(15) and scores for 64 different cell types using the xCell algorithm(16), separately for both the DC and VC biopsies. We identified significant (FDR<0.05) associations between the cell type scores and Module 9 that were in common between the DC and VC biopsies (**Fig. S6)** and identified 8 cell types (via xCell) including dendritic cells, activated dendritic cells, plasmacytoid dendritic cells, macrophages, M1 macrophages as well as CD8+ effector memory T cells, CD8+ central memory T cells, and T regulatory cells (**Fig. 4C).** Taken together, the progressive/persistent biopsies in the Proliferative subtype have down-regulated expression of Module 9 compared with regressive biopsies that correlates with reduced signals from both innate and adaptive immune cell populations.

### Immunofluorescence reveals progression-associated modulation of macrophages and T cells in Proliferative PMLs

In order to confirm the relationship between the immune cell types associated with Module 9 and histologic progression/persistence of PMLs in the *Proliferative* subtype, immunofluorescent staining of macrophages/monocytes (n=52 regions enumerated from n=16 subjects), CD4 (n=50 regions enumerated from n=17 subjects), and CD8 T cells (n=47 regions enumerated from n=16 subjects) was performed (**Table S5**). The results were analyzed across all subjects assayed within the *Proliferative* subtype and across the subset of subjects where the lesion outcome (progression/persistence versus regression) was concordant with the Module 9 GSVA score (denoted as concordant set). Staining of CD68, a pan macrophage (and tumor associated macrophage) marker, suggestive of M1 type macrophages, was increased in progressive/persistent lesions (p<<0.001 in the concordant set). In contrast, staining of CD163 in combination with CD68, thought to be suggestive of M2 type macrophages, were decreased among the progressive/persistent lesions in the *Proliferative* subtype (p<<0.001 using all subjects and p=0.0007 in the concordant set, respectively) (**Fig. 4D-E**). Additionally, CD4 T cells were increased (p<<0.001 in the concordant set) and CD8 T cells were decreased (p<<0.001 in the concordant set) in PMLs that progress/persist. Interestingly, among progressive/persistent lesions, the CD8 T cells had a distinct localization pattern (p=0.07 in the concordant set), where CD8 T cells both lined and were embedded within the epithelium in areas where dysplasia is present (**Fig. 4D**). The immunofluorescence results did not reach significance, with the exception of CD163, when just the lesion outcome was used without regard to the Module 9 score.

## Discussion

Lung squamous cell carcinoma (LUSC) is the second most common form of lung cancer and arises in the epithelial layer of the bronchial airways. It is often preceded by the development of lung squamous premalignant lesions (PMLs). The presence of dysplastic persistent and or progressive PMLs is a marker of increased risk for LUSC*(6)*. Currently, however, we lack effective tools to identify PMLs at highest risk of progression to invasive carcinoma*(7)*. The development of markers predictive of disease progression will be important in identifying patients at highest risk for LUSC development and in identifying biological pathways exploitable for LUSC chemoprevention. Towards this goal, we profile via RNA-Seq bronchial brushes and endobronchial biopsies obtained from subjects undergoing longitudinal lung cancer screening by chest computed tomography (CT) and autofluorescence bronchoscopy. We identify four transcriptionally distinct groups of biopsies, one of these we label *Proliferative* and find it to be associated with high-grade dysplasia. Patients with *Proliferative* PMLs can also be identified via gene expression measured from cells in the non-involved large airway epithelium. We further find that persistent/progressive *Proliferative* PMLs are characterized by decreased expression of genes involved in interferon signaling and antigen processing/presentation pathways. Consistent with these gene expression findings we find that progressive/persistent *Proliferative* PMLs are depleted for CD68+/CD163+ macrophages and CD8 T cells by immunofluorescence. Collectively, these data suggest both the potential to identify a subset of patients with progressive/persistent LUSC PMLs, who are at risk for developing invasive lung cancer, on the basis of airway gene expression; as well as the potential to decrease the risk for progression in these patients by augmenting the immune response associated with regression.

Previous studies indicate a range of genomic alterations associated with bronchial dysplasia. Increased expression of *EGFR* and Ki67 staining of epithelial cells is associated with increasing histologic severity and subsequent histologic progression*(6, 17)*. Altered protein levels of *TP53, CCND1, CCNE1, BAX, and BCL2* have been associated with *CIS* or lung cancer occurrence independent of histological grade(18). Telomere shortening and maintenance(19) and loss of heterozygosity in regions frequently detected in lung cancer (3p, 5q, 9p, 13q, 17p) have been observed in early hyperplasia/metaplasia lesions(20-22) and found to increase in frequency and size in higher-grade dysplasia. Genomic gains in loci containing *SOX2, TP63, EGFR, MYC, CEP3*, and *CEP5* are also associated with progression of high-grade dysplasia*(23)*. Despite the numerous genomic alterations associated with PML histological grade and progression, we lack a comprehensive PML molecular classification system to complement the pathologic classification of PML. We pursued an unsupervised class discovery approach that led to the identification of four distinct molecular PML subtypes (*Proliferative, Inflammatory, Secretory*, and *Normal*). The transcriptional patterns differentiating the PML subtypes are robust and a 22-gene panel identified in the Discovery Cohort can be used to distinguish between the different molecular subtypes in an independent Validation Cohort. The *Proliferative* subtype is enriched with dysplastic PMLs from current smokers and is characterized by up-regulation of metabolic (OXPHOS/ETC/TCA) and cell cycle pathways and down-regulation of cilia-associated pathways. Previous work indicates increases in metabolic pathways in the airways of subjects with dysplastic lesions(13), in PMLs adjacent to LUSC tumor*(24)*, and in smokers at high-risk for lung cancer*(25)* as well as increases in proliferation (via Ki67 levels, as mentioned above) that have been utilized as an endpoint in lung cancer chemoprevention*(26, 27)*. Identification of patients with *Proliferative* lesions may be useful to enrich lung cancer chemoprevention trials with high-risk subjects or to identify patients who would benefit from more frequent lung cancer screening. The *Inflammatory* subtype is predominated by PMLs from former smokers, but interestingly is not significantly enriched for dysplasia, despite similarly decreased expression of cilia-associated pathways, suggesting an abnormal epithelium. The *Inflammatory* subtype also shows increased expression of a gene module enriched for genes involved in inflammation and regulation of lymphocytes and leukocytes (Module 8). This gene module is also elevated in *Secretory* lesions predominated by lesions from current smokers and exhibiting increased expression of goblet cell markers. Interestingly, *IL1B* is part of this inflammation-related gene module, which is of great interest as the inhibition of *IL1B* has recently been shown to reduce lung cancer incidence*(28)*.

Our prior work has extensively studied gene expression alterations in normal-appearing airway epithelium by profiling cells obtained via brushing the mainstem bronchus during bronchoscopy*(8, 29-35)*. As part of this work, we have described gene expression alterations that reflect the presence of bronchial dysplasia*(31)*. In the current study, for the first time we have both bronchial brushes and endobronchial biopsies collected during the same procedure allowing us to identify gene expression differences in bronchial brushings from normal appearing airway which indicate the presence of *Proliferative* subtype PMLs. In both the Discovery and Validation cohorts, applying the predictor used to identify *Proliferative* subtype PMLs (based on PML biopsy gene expression) to the gene expression data from the normal-appearing airway brushings resulted in predictions of the *Proliferative* subtype that were very specific (91%) but not sensitive (31-38%). Brushes classified as *Proliferative* have increased expression of cell cycle pathways and decreased expression of cilia-associated genes, suggesting that they are more similar to squamous metaplasia than normal epithelium. Potentially, a subset of patients may harbor widespread airway damage that serves as a marker for the presence of this type of high-grade PML leading to modest sensitivity, but high specificity. In other cases, the area of damage that gives rise to these *Proliferative* PMLs may be more localized, and therefore potentially more difficult to detect by brushing contributing to decreased sensitivity. These findings suggest that therapeutics to target changes throughout the entire airway epithelium may be necessary in some subjects, whereas, more site-specific ablation (e.g. photodynamic therapy) may be more effective in certain cases. Another possibility and area of future research, is that a *Proliferative* subtype brush is a predictor of incident LUSC.

The molecular profiling of PMLs and the identification of gene co-expression modules also provides an opportunity to identify the molecular determinants of subsequent PML progression. One of the nine gene co-expression modules used to define the molecular subtypes was significantly different between biopsies that progress or persist compared to biopsies that regress within the *Proliferative* subtype in both the DC and VC cohorts. The module contains genes whose expression is decreased in the persistent/progressive biopsies that are involved in interferon signaling and antigen processing and presentation. These gene expression changes were correlated with a decreased abundance of innate and adaptive immune cells via computational prediction. By immunofluorescent staining of FFPE biopsy sections we confirmed that the progressive/persistent *Proliferative* lesions with low Module 9 GSVA scores had fewer CD163+ macrophages and CD8+T cells and the CD8+T cells had a distinct localization pattern. These lesions also contained greater numbers of CD4+T cells, and it will be important in future work to assess if these cells are T regulatory cells promoting an immune suppressive environment.

The presence of tumor-associated macrophages with the polarized phenotypes (M1 as pro-inflammatory or M2 as anti-inflammatory) has been associated with lung cancer prognosis. The presence of predominantly M2 macrophages, marked by the expression of CD163, has been associated with worse survival. However, in the context of lung PMLs this relationship is not well studied. Our finding that regressive *Proliferative* PMLs have more CD163+ cells and increased expression of genes involved in IFNg signaling is consistent with what has been seen in the PMLs that precede oral squamous cell carcinoma where the presence of CD163+ macrophages with active IFNg signaling is associated with better outcomes*(36)*. Additionally, we observed fewer CD8+ T cells and lower expression of HLA class I genes and B2M in progressive/persistent lesions within the *Proliferative* subtype. Disruptions in proper T cell mediated immunosurveillence have been described in several studies showing that impaired HLA class I antigen processing and presentation including down-regulation or loss of B2M*(37, 38)* and interferon signaling*(39)* in lung tumors affects response and acquired resistance to checkpoint inhibitors. Lung tumors lacking an HLA-I complex had lower cytotoxic CD8+ lymphocyte infiltration, and this was also associated with lower levels of PD-L1. Additionally, studies have also suggested negative impacts on efficacy of check point inhibitors as well as survival in patients with LC that have tumors with increased CD4+ T cells expressing T regulatory markers (FOXP3, CD25) resulting in immunosuppressive state suggested to hinder the recruitment and effector functions of CD8+ T cells*(40, 41)*. Future DNA sequencing data on the PMLs profiled here may indicate heterozygous or homozygous loss of B2M or mutations in other genes in the interferon and antigen processing and presentation pathways; however, even in the case of acquired resistance, mutations and copy number changes could not explain the down-regulation of these pathways across all subjects, suggesting that other epigenetic alterations or signaling pathways may play a role. In fact, epigenetic therapy, specifically DNA methyltransferase inhibitors*(42)*, has been shown to enhance response to immune checkpoint therapy and up-regulate many of the genes down-regulated in progressive/persistent lesions within the *Proliferative* subtype including HLA class I genes (*HLA-B* and *HLA-C*), *B2M, CD58, TAP1*, immune-proteasome subunits *PSMB9* and *PSMB8*, and the transcription factor *IRF9.* Unraveling the mechanisms of innate and adaptive immune down-regulation in this subset of PMLs will be important to identifying potential immunoprevention therapies.

Our data suggests that there are subtype-specific transcriptomic alterations predictive of subsequent LUSC premalignant lesion progression that are the result of a lack of infiltrating immune cells in the lesion microenvironment. These data suggest that biomarkers for determining PML subtype and assessing immune infiltration may have utility for the detection of aggressive PMLs that require more intensive clinical management and genes altered in these PMLs may serve as lung chemoprevention candidates. These biomarkers could either be measured directly in PML tissue, or our data also suggests the potential that they could be measured in a surrogate tissue such as bronchial airway epithelium. A benefit of biomarkers predicting aggressive PML behavior measured in surrogate tissue is the potential that these biomarkers might also predict the behavior of PMLs not directly observed during bronchoscopy. Future studies are needed to address the specific mechanism of impaired immunosurveillence in progressive/persistent lesions in the *Proliferative* subtype including high coverage DNA sequencing, characterization of neoepitope burden, assessment of epigenetic alterations, and comprehensive characterization of the immune populations identified.

## Materials and Methods

### Subject Population and Sample Collection

Endobronchial biopsies and brushings were obtained from high-risk subjects undergoing lung cancer screening at approximately 1-year intervals by white light and auto-fluorescence bronchoscopy and computed tomography at Roswell. The bronchoscopy included visualization of the vocal cords, trachea, main carina, and orifices of the sub-segmental bronchi visible without causing trauma to the bronchial wall. All abnormal and suspicious areas are biopsied twice and the lung anatomic location is recorded (**Fig. S7, Table S7**). One biopsy was used for routine pathological evaluation and the other for molecular profiling. Additionally, a brushing was obtained from a normal appearing area of the left or right mainstem bronchus for research. Morphological criteria used to evaluate the biopsies are in accordance with World Health Organization (WHO) guidance*(43)*. Eligibility for screening includes either a previous history of aerodigestive cancer and no disease at the time of enrollment or age greater than 50, a current or previous history of smoking for a minimum exposure of 20 pack-years and at least one additional risk factor including moderate chronic obstructive pulmonary disease (COPD) (defined as forced expiratory volume (FEV1) < 70%), confirmed asbestos related lung disease or a strong family history of lung cancer (at least 1-2 first degree relatives). All research specimens were stored in RNA Allprotect (Qiagen) and stored at −80 degrees C.

Subjects were selected that had biopsies collected in repeat locations via serial bronchoscopies; however, after RNA isolation, samples from 3 subjects had a single biopsy and 1 subject had a single brushing. mRNA sequencing was performed on a discovery cohort (DC) of samples comprising of endobronchial biopsies and brushes collected between 2010 and 2012 (n=30 subjects, n=197 biopsies, and n=91 brushings). mRNA sequencing was subsequently performed on a validation cohort (VC) of samples comprising of endobronchial biopsies and brushes collected between 2012 and 2015 (n=20 subjects, n=111 biopsies, and n=49 brushings). Brush histology was defined by the worst biopsy histology observed at the same time point. Biopsy progression/regression was defined for each biopsy based on the histology of the biopsy and the worst histology recorded for the same lung anatomic location in the future. Histology changes between normal, hyperplasia, and metaplasia were classified as “normal stable”, decreases in histological dysplasia grade or changes from dysplastic histology to normal/hyperplasia/metaplasia were classified as “regressive”, lack of future histological data was classified as “unknown”, and everything else was classified as “progressive/persistent.” The Institutional Review Boards at Boston University Medical Center and Roswell approved the study and all subjects provided written informed consent.

### RNA-Seq library preparation, sequencing, and data processing

Total RNA was extracted from endobronchial biopsies and bronchial brushings using miRNeasy Mini Kit or AllPrep DNA/RNA/miRNA Universal Kit (Qiagen). Sequencing libraries were prepared from total RNA samples using Illumina TruSeq RNA Kit v2 and multiplexed in groups of four using Illumina TruSeq Paired-End Cluster Kit. Each sample was sequenced on the Illumina HiSeq 2500 to generate paired-end 100-nucleotide reads. Demultiplexing and creation of FASTQ files were performed using Illumina CASAVA 1.8.2 or BaseSpace. Samples were aligned using hg19 and 2-pass STAR*(44)* alignment. Gene and transcript level counts were calculated using RSEM*(45)* using Ensembl v74 annotation. Quality metrics were calculated by STAR and RSeQC*(46)*. Samples were excluded were sex annotation did not correlate with gene expression across *CYorf15A* (ENSG00000131002), *DDX3Y* (ENSG00000067048), *KDM5D* (ENSG00000012817), *RPS4Y1* (ENSG00000129824), *USP9Y* (ENSG00000114374), and *UTY* (ENSG00000183878) (n=4 samples). Sample relatedness within a patient was confirmed using Peddy software*(47)*. Samples with a high-rate of heterozygosity (more than 3 standard deviations above the median) or samples with low relatedness to samples from the same patient (more than 3 standard deviations below the median) were removed from further analyses (n=11 samples, 2 brushes and 9 biopsies). Samples were subsequently divided into the discovery and validation cohorts (as outlined above) and by tissue type (biopsy or brush). Subsequent sample and gene filtering was conducted separately on each set as follows: First, EdgeR*(48)* was used to compute normalized data (library sizes normalized using TMM, trimmed mean of M-values, and log2 counts per million computed) and genes were excluded that either had an interquartile range equal to zero or a sum across samples equal or less than 1. Samples were excluded based on values greater than 2 standard deviations from the mean for more than one of the following criteria: 1) mean Pearson correlation with all other samples calculated across all filtered genes 2) the 1^st^ or 2^nd^ principal components calculated using the filtered gene expression matrix 3) transcript integrity number (TIN, computed by RSeQC). After sample filtering, gene filtering was recomputed as described above on the final set of high-quality samples. The data are available from NCBI’s Gene Expression Omnibus using the accession GSE109743.

### Derivation of molecular subtypes

The DC biopsies (n=190 samples, n=16653 genes) and brushes (n=89 samples, n=16058 genes) were used to derive the molecular subtypes. Two additional RNA-Seq datasets were used during the derivation of the molecular subtypes: the TCGA squamous cell carcinoma (LUSC) tumors(10) (n=471 samples, n=17887 genes) and a dataset of tracheobronchial samples from mice treated with n-nitrosotris-(2-choroethyl)urea (NTCU) (n=25 samples, n=14897 genes). The mice develop lesions that are histologically and molecularly comparable to human lesions and that progress to LUSC and the samples represent a range of histology (normal, mild dysplasia, moderate dysplasia, severe dysplasia, carcinoma in situ (CIS), and LUSC tumor) (**Supplementary Materials and Methods**). The mouse data are available from NCBI’s Gene Expression Omnibus using the accession ID GSE111091. Sample and gene filtering from the TCGA LUSC tumors and the mouse tissue were processed as described in the **Supplementary Materials and Methods**.

Weighted correlation network analysis(9) (WGCNA) was used with default parameters to derive Modules of gene co-expression across the 4 datasets described above. Residual gene expression values adjusting for RNA quality (median TIN) and batch (Illumina flow cell) were used as input for WGCNA for the biopsy and brush datasets. For the mouse dataset, residual gene expression values adjusting for RNA quality (median TIN), mouse strain, and sample type (laser capture microdissected versus whole tissue) were used as input for WGCNA. Log2 counts per million (cpm) values were used as input for WGCNA for the LUSC tumor samples. Gene sets were created for each co-expression Module for each dataset and then combined to create a compendium of gene sets generated from each of the 4 datasets. For each gene set in the compendium, the first principal component (PC1) was calculated across each z-score normalized dataset. For each dataset, a Pearson correlation matrix of PC1 values across all gene sets in the compendium was computed and thresholds were set as follows: r>0.85 was set to 1 and r<=0.85 set to 0. The four matrices were subsequently summed, and gene sets derived from biopsy co-expression Modules that were correlated to another non-biopsy derived gene set across all datasets were retained (n=9 Modules retained). The genes defining the retained biopsy Modules were required to be present in the biopsy Module and at least in one of the correlated gene sets.

The filtering process above yielded a reduced set of genes (n=3,936) that was used to define the molecular subtypes in the biopsy data. The residual gene expression values across the reduced set of genes for the discovery biopsies was used as input for consensus clustering*(49)*. Consensus clustering was performed setting k (number of groups) to 10, the number of iterations to 1000, the subsampling to 80%, the clustering algorithm to partitioning around mediods, and the distance metric to Pearson correlation. The optimal value for k was 4 based on the relative change in area under the cumulative distribution function calculated based on the consensus matrix for each k.

### Molecular subtype predictor

The DC biopsies across the filtered genes were used to derive a molecular subtype predictor. First, Pearson correlation metrics were determined between each gene and the Module eigengenes (PC1 for each of the 9 Modules). Genes were retained as part of a Module if the correlation value was the highest for the Module in which it was assigned. The average Pearson correlation of the retained genes to the Module eigengene was computed, and the number of genes chosen from each Module for the predictor was inversely proportional to this metric. Second, the genes most highly correlated to the Module eigengene were chosen to represent the Module in the predictor. The 22 genes resulting from this analysis across the DC biopsy data were used to train a nearest centroid predictor using the pamr package with a threshold of zero and predict the molecular subtype across the VC biopsies. Prior to predicting the molecular subtype of these test sets, the training and test sets were combat*(50)* adjusted and z-score normalized across combined training and test data. Using the methods described above we derived molecular subtypes using consensus clustering across the VC biopsies and compared these to the predicted subtypes.

### Identification of biological processes associated with gene modules and molecular subtypes

Biological processes and pathways enriched in each of the nine Modules used to discover the molecular subtypes in the DC were identified using EnrichR*(51)*. Each Module was separated into genes positively or negatively correlated with the Module eigengene, the Ensembl IDs were converted to Gene Symbols using biomaRt, and the following databases were queried: GO Biological Process 2015, KEGG 2016, WikiPathways 2016, TargetScan microRNA, Transcription Factor PPIs, TRANSFAC and JASPAR PWMs, OMIM Disease, Reactome 2016, and Biocarta 2016. Processes/pathways with an FDR<0.05 were considered to be significantly enriched. The contribution of each gene Module to the DC biopsy molecular subtypes was evaluated by testing if GSVA(14) scores for each Module were significantly (FDR<0.05) associated with the molecular subtypes using a linear mixed effect model with patient as a random effect via limma.

### Identification of clinical and biological phenotype associations with molecular subtype

The molecular subtypes in the DC biopsies were annotated according to the behavior of each gene Module by calculating whether or not GSVA(14) scores for each Module were significantly up- or down-regulated (FDR<0.05) in a particular molecular subtype versus all other samples using a linear mixed effects model with patient as a random effect via limma. Additionally, the biological pathways and transcription factors associated with each subtype were identified using GSEA*(52)* and mSigDB*(53)* gene sets using genes ranked by the t-statistic for their association with each subtype. The ranked lists were created using the limma*(54)* and edgeR*(48)* packages to identify differentially expressed genes associated with subtype membership. Each linear model used voom-transformed*(55)* data and included membership in the subtype of interest, batch, and RNA quality (TIN) as covariates and patient as a random effect. Pathways enriched in the ranked lists (FDR<0.05) were used to annotate the molecular subtypes. FDR values for individual genes were derived from this analysis or analogous models using only samples of normal/hyperplasia histology or dysplasia histology.

For the DC and VC biopsies, residual gene expression values were used to predict smoking status, LUSC tumor subtype, and the relative abundance of epithelial and immune cells for each sample. Smoking status (current versus former/never) was predicted for each sample as described previously(13). Smoking status was determined at each time point for each subject by calculating the mean of the prediction scores (>0 for current prediction and <0 for former/never prediction) across all biopsies and brushes sampled. The LUSC tumor subtype was determined as described previously(11) across the genes predictive of the LUSC molecular subtype(12). The ESTIMATE algorithm*(56)* was used to infer relative epithelial, stromal, and immune cell content. Immune cell type specific signatures from Bindea *et al*.(15) and epithelial cell type specific signatures from Dvorak *et al.*(50) were used to generate GSVA(14) scores across samples for each signature. Additionally, residual gene expression values calculated using log RPKM values were inputted into the xCell(16) to infer relative abundances of 64 different cell types. The above categorical phenotypes along with additional clinical variables such as biopsy histology, subject, previous lung cancer history, sex, and biopsy progression/regression status were associated with molecular subtype using Fisher’s Exact Test. Continuous variables were associated with molecular subtype using a linear model via limma.

### Relationship between the biopsies and brushes

We wanted to quantify the predictive performance of the brush with regards to the presence of a biopsy of the Proliferative subtype. A subset of the 22-gene molecular subtype predictor was used to predict the presence or absence of the Proliferative subtype across the DC and VC brushes and biopsies. Specifically, we used 8 genes (out of the 22) that corresponded to Modules 4 through 7 (significantly up- or down-regulated in the Proliferative subtype) to classify samples as Proliferative or not using the same methodology described above for the molecular subtype predictor. Sensitivity and specificity performance metrics were calculated based on the ability of a Proliferative subtype prediction in the DC or VC brushes to indicate the presence of at least one biopsy of the Proliferative subtype. In order to further understand the Proliferative subtype predictions in the brushes, we analyzed the behavior of the modules that define the Proliferative subtype in the DC biopsies (based on methods above) across the DC and VC brushes.

### Immunofluorescent staining and quantitation

Standard formalin fixation and embedding techniques were employed at Roswell where 5-micron sections were cut from the FFPE samples used for the routine pathological evaluation at Roswell (**Table S5**). Prior to staining, samples were de-waxed with xylene and rehydrate through a graded series of ethanol solutions. AR or citrate buffer was used for antigen retrieval, tissue was incubated with primary antibodies overnight at 4°C and probed with secondary antibodies with fluorescent conjugates (Invitrogen Alexa Fluor 488, 594, 647) for 1 hour at room temperature. Immunostaining was performed using the primary antibodies listed in **Table S8**. Imaging was performed using an Aperio Slide Scanner for scoring and a Carl Zeiss Axio (20x and 40 x objectives) and a Carl Zeiss LSM 710 NLO confocal microscope for capturing additional images. Digital slides were analyzed with the Definiens Tissue Studio (Definiens Inc.) for the enumeration of immunofluorescence staining. The enumeration of the immunofluorescence scored each stain including DAPI positive cells. The enumeration was conducted on different regions (independent areas of tissue) present on a slide (1-5 regions/biopsy) for each biopsy. For each region, the percentage of positively staining cells for a given protein was calculated by dividing the number of positively stained cells by the total number of DAPI positive cells. A binomial mixed effects model via the lme4 R package was used to assess differences in the percentages of cells staining positive for a given protein in each region between progressive/persistent versus regressive biopsies using the total cells stained in each region as weights and adjusting for the slide number as a random effect. The models were used across samples from the *Proliferative* subtype and across samples from the Proliferative subtype where the biopsy outcome (progressive/persistent versus regressive) agreed with the Module 9 GSVA score (scores less than 0 are associated with progression/persistence and scores greater than 0 are associated with regression). Each region was also qualitatively scored as either positive or negative for having a distinct CD8 T cell localization pattern where cells lined and were embedded within the epithelium.

## Supplementary Materials

Supplementary Materials and Methods.

Fig. S1. Distribution of Molecular Subtypes by Subject.

Fig. S2. Immunofluorescent Staining Quantitation of Proliferation, Basal Cell, and Ciliated Cell Markers across the Molecular Subtypes.

Fig. S3. Boxplots of Select Genes and Cell Type Deconvolution Results across the Discovery and Validation Cohorts by Molecular Subtype.

Fig. S4. Heatmap of the 22-gene Molecular Subtype Classifier in the Discovery and Validation Cohort Biopsies.

Fig. S5. Gene module behavior across the Molecular Subtypes in the Discovery and Validation Cohort Biopsies.

Fig. S6. Concordance between Module 9 and two Cell Type Deconvolution Analyses. Fig. S7. Tracheobronchial Map.

Table S1. Batch Information and Alignment Statistics on Samples in both the Discovery and Validation cohorts.

Table S2. Summary of Gene Modules.

Table S3. Pathways enriched in the Gene Modules.

Table S4. Molecular Subtype associations with Clinical and Biological Characteristics within the Discovery Cohort (DC) and the Validation Cohort (VC).

Table S5. List of Samples used for Immunofluorescence Studies.

Table S6. Statistical associations between Progression/Persistence versus Regression within each Molecular Subtype and Cohort (DC and VC) for each Gene Module.

Table S7. Lung sites where Endobronchial Biopsies were obtained. The site code, name, Table S8. Antibodies used in the Immunofluorescence Studies.

Data file S1. Complete list of pathways enriched in gene modules.

## Author Contributions

Study conception and design (JB, SAM, JDC, CP, CM, MS, MER, MEL, AES, SJP, CS); collection of clinical samples (SSD); study coordination (JV); sample processing and library preparation (GL, SZ, HL); data analyses (JB); immunofluorescence (SAM, GD, KK, SMD, JB); writing of the manuscript (JB, SAM); editing of the manuscript (JB, SAM, JDC, CM, MEL, AES, JV).

## Data and materials availability

GSE109743

